# Predicting unrecognized enhancer-mediated genome topology by an ensemble machine learning model

**DOI:** 10.1101/2020.04.10.036145

**Authors:** Li Tang, Matthew C. Hill, Jun Wang, Jianxin Wang, James F. Martin, Min Li

## Abstract

Transcriptional enhancers commonly work over long genomic distances to precisely regulate spatiotemporal gene expression patterns. Dissecting the promoters physically contacted by these distal regulatory elements is essential for understanding developmental processes as well as the role of disease-associated risk variants. Modern proximity-ligation assays, like HiChIP and ChIA-PET, facilitate the accurate identification of long-range contacts between enhancers and promoters. However, these assays are technically challenging, expensive, and time-consuming, making it difficult to investigate enhancer topologies, especially in uncharacterized cell types. To overcome these shortcomings, we therefore designed LoopPredictor, an ensemble machine learning model, to predict genome topology for cell types which lack long-range contact maps. To enrich for functional enhancer-promoter loops over common structural genomic contacts, we trained LoopPredictor with both H3K27ac and YY1 HiChIP data. What’s more, the integration of several related multi-omics features facilitated identifying and annotating the predicted loops. LoopPredictor is able to efficiently identify cell type-specific enhancer mediated loops, and promoter-promoter interactions, with a modest feature input requirement. Comparable to experimentally generated H3K27ac HiChIP data, we found that LoopPredictor was able to identify functional enhancer loops. Furthermore, to explore the cross-species prediction capability of LoopPredictor, we fed mouse multi-omics features into a model trained on human data and found that the predicted enhancer loops outputs were highly conserved. LoopPredictor enables the dissection of cell type-specific long-range gene regulation, and can accelerate the identification of distal disease-associated risk variants.

## INTRODUCTION

Developmental gene regulatory networks rely on cis regulatory elements, like enhancers, to drive gene expression patterns in both space and time in a cell type-specific fashion. Enhancer evolution also plays an important role in driving morphological divergence (1). Moreover, enhancers play a role in maintaining cell identity and responding to external stimuli, like injury and infection. Many enhancers work over long genomic distances through the formation of topological loops to promoters to regulate gene expression (2). Importantly, the majority of identified disease associated genetic variants uncovered through genome-wide association studies (GWAS) reside in non-coding intergenic regions that often can be ascribed enhancer activity. Hence, identifying the promoters looped to these variants in a cell type-specific manner is important for determining their pathological roles (3–5).

In the past decade, high-throughput based Chromosome Conformation Capture (3C) techniques have been developed to understand genome architecture (6). High-throughput Chromosome conformation capture (HiC) (7) identifies physical genomic interactions in a genome-wide fashion, but requires deep sequencing to achieve high resolution, which is costly and difficult to apply on a large-scale. Chromatin Interactive Analysis by Paired-End Tag Sequencing (ChIA-PET) aims to detect the specific long-range interactions associated with a protein of interest (8). However, ChIA-PET requires a large number of cells as input (8). Recently, HiChIP, a protein-centric chromatin conformation method was developed, which requires lower input and also achieves a larger number of conformation-informative reads compared to traditional ChIA-PET protocols (9). HiChIP has been used to produce contact data for a number of key chromatin binding factors, including YY1, and cohesion (9–11). H3K27ac, an active enhancer- and promoter-associated histone mark, distinguishes active enhancers from inactivate enhancers (12–14). And H3K27ac HiChIP data identifies high-confidence functional enhancer-promoter interactions (10). Similarly, YY1 binds to active enhancers and promoter–proximal elements, and acts as a structural regulator of enhancer-promoter interactions to facilitate gene expression, making it a suitable marker for identifying distal acting enhancer-promoter pairs (11). However, HiChIP like other 3C methodologies still requires specialized reagents, equipment, and high depth sequencing making it difficult to perform on a large scale.

We therefore constructed an ensemble machine learning model, LoopPredictor, to predict enhancer mediated loops in a genome-wide fashion across different cell lines and species. H3K27ac/YY1 HiChIP data in K562, GM12878, and HCT116 cell lines as well as other multi-omics features (e.g. ChIP-Seq, RNA-seq, ATAC-seq) were integrated to train the model. Importantly, the predicted genomic contacts output from our pipeline were consistent with published functionally validated gene-enhancer loops. Finally, when non-topological multi-omics features from a murine cell line were gathered and fed into the model, which was trained on human data sets, the cross-species prediction overlapped favorably with H3K27ac HiChIP performed on the mouse cell line. LoopPredictor leverages the long-range enhancer mediated loop enrichment derived from H3K27ac/YY1 HiChIP datasets, and then integrates user-provided multi-omics feature inputs to predict active enhancer mediated loops for any cell type with high sensitivity.

## RESULTS

### Identifying active enhancer-promoter loops with H3K27ac and YY1 HiChIP

Previous studies have reported that H3K27ac HiChIP identifies high-confidence active enhancer-promoter chromatin loops (10). To assess the quality of this method, we analyzed H3K27ac HiChIP loops from K562 cells, a human bone marrow-derived cell line, and found that the majority of loops were enhancer mediated, with a small percentage of promoter-promoter interactions (**Fig. 1A)**. Super enhancers play critical roles in many important biology events such as cell identity, development, and oncogenesis (15–17). We analyzed the cis regulatory elements with super enhancer status from H3K27ac HiChIP anchors and found that super enhancers account for 5.6% of all enhancers (**Fig. 1B**). Gene ontology (GO) analysis of these super enhancer anchors indicated that they contribute to the cell identity of K562 cells, confirming the suitable quality of this dataset for assaying biologically relevant topological interactions (**Fig. 1C**). To determine the transcription factor composition of H3K27ac enhancer-promoter loops in an unbiased fashion, we carried out motif analysis on H3K27ac HiChIP loop anchors (**Fig. 1D**). The results indicated that the YY1 motif is significantly enriched (−log2(p-value) < 100) in H3K27ac loop anchors, and is also highly expressed at the transcriptional level in K562 cells. Given that YY1 has been suggested as a faithful marker of active enhancers and proximal promoters (11), we hypothesized that H3K27ac and YY1 co-occupied enhancer-promoter loops should overlap favorably and represent a high-confidence set of active distal enhancers (**Fig. 1E**). To verify this hypothesis, we compared H3K27ac HiChIP loops with YY1 HiChIP loops. We found the majority of YY1 HiChIP loops (86 %) overlapped with H2K27ac loops. Importantly, the overlapping set of distal interactions found were primarily enhancer-mediated loops (**Fig. 1F**). For example, the *CLIP2* locus showed similar H3K27ac and YY1 ChIP-seq profiles with both H2K27ac and YY1 enriched topological interactions present between distal elements and the promoter (**Fig. 1G**). Together, these results indicated that H3K27ac and YY1 HiChIP data could be combined to comprehensively characterize the full suite of highly active enhancer-promoter pairs present in a cell type of interest. The experimental enrichment of active enhancer loops over structural loops (e.g. CTCF-mediated) commonly associated with HiC data, makes H3K27ac and YY1 HiChIP ideal for use in the construction of a novel machine learning model to predict unrecognized active enhancer-mediated loops.

**Figure 1.**
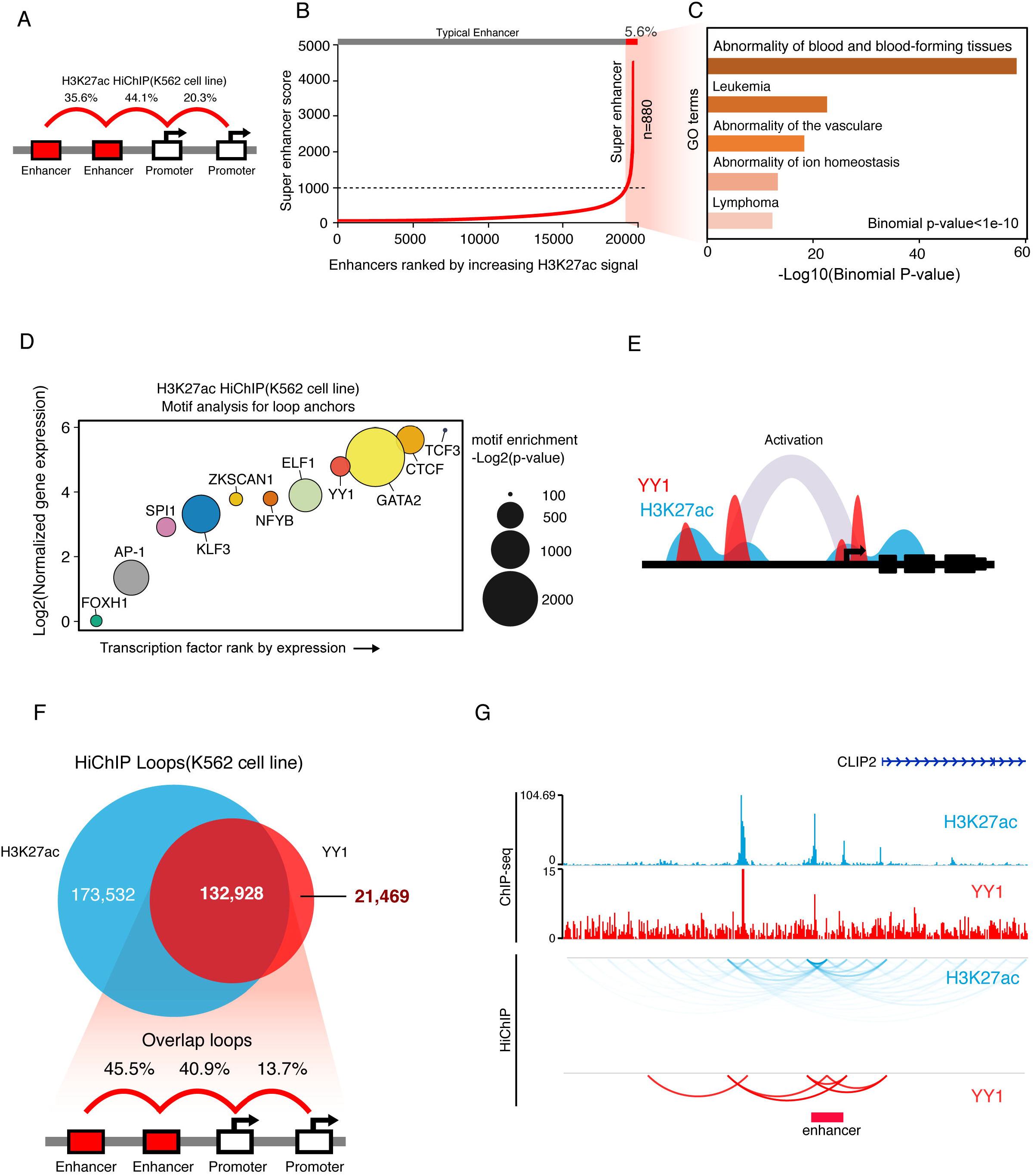
H3K27ac and YY1 HiChIP demarcate active enhancer loops. (A) Proportion of annotated loop types for K562-H3K27ac HiChIP data. Each loop identified with an FDR < 0.05, and pair-end tag number ≥ 2. ChromHMM was used to annotate the anchors, with only enhancer and promoter type anchors being retained. Majority of loops are enhancer mediated (79.7%). (B) Super enhancer plot for anchors in K562-H3K27ac HiChIP data. Slope threshold was set at 1000. A total of 880 super enhancers were found, which accounted for 5.6% of all enhancers. Super enhancer signal derived from H2K27ac ChIP-seq data. (C) GO analysis for super enhancer anchors in K562-H3K27ac HiChIP data (binomial p-value less than 1 e^−10^). (D) *De novo* motif enrichment analysis on K562-H2K27ac loops. Transcription factors ranked by normalized gene expression. The size of each point indicates the motif enrichment *P* value. The transcription factors with high motif enrichment (–log2(p-value) > 100) and gene expression are shown. The YY1 motif was significantly enriched (−log2(p-value) = 306). (E) Diagram depicting the putative co-enrichment of H3K27ac and YY1. (F) Top, Venn diagram showing the intersect of K562-YY1 and K562-H3K27ac HiChIP datasets. Bottom, proportion of each annotated loop category from the overlapping H3K37ac and YY1 HiChIP loops (n = 132,928). The majority of the overlapping loops were enhancer associated (86.4%). (G) Genome browser tracks showing H3K27ac and YY1 ChIP-seq signals and topological interactions.

### An ensemble machine learning model to predict enhancer mediated loops

To overcome the shortcomings associated with large-scale HiChIP, HiC, and ChIA-PET experiments, we developed an ensemble machine learning model, named LoopPredictor to predict enhancer mediated loops from multi-omics features (**Fig. 2A**). First, training datasets were gathered, which could be separated into features and targets. LoopPredictor integrated multi-omics datasets, including ChIP-seq/CUT&RUN (3,18–22), ATAC-seq (18,23–25), RNA-seq (3,26), and Reduced Representation Bisulfite Sequencing (RRBS) (3) as features, while also implementing four published HiChIP datasets as targets for the prediction, including K562-YY1 (11), K562-H3K27ac (10), HCT116-YY1 (11), GM12878-H3K27ac (10). Second, the algorithm core of LoopPredictor was constructed, which consists of two components: ATP and CP.

**Figure 2.**
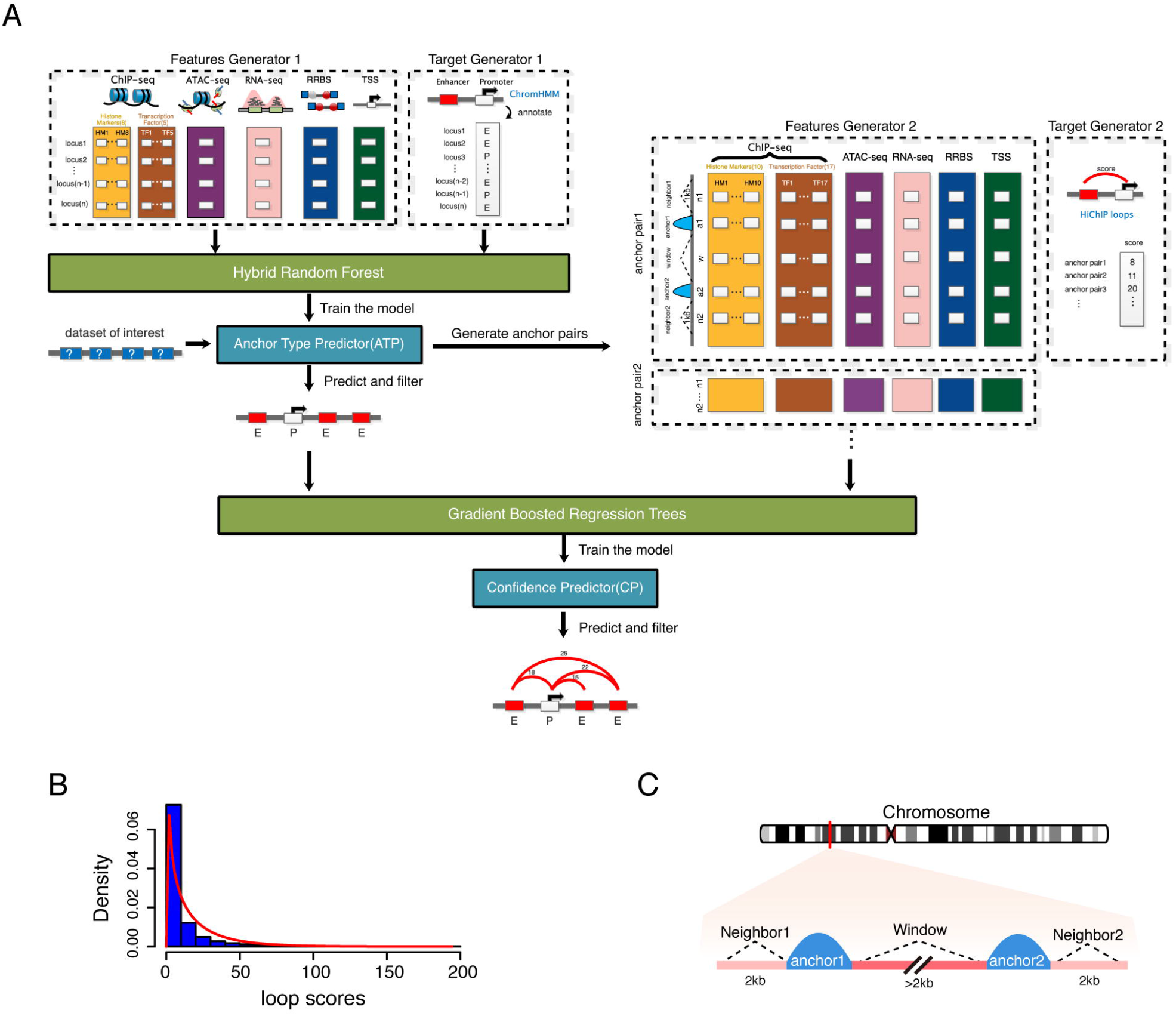
LoopPredictor, an ensemble machine learning model. (A) The LoopPredictor algorithm. H3K27ac and YY1 HiChIP datasets and multi-omics features (e.g. ChIP-Seq, RNA-seq, ATAC-seq, and RRBS) were first processed and then integrated to train the model. Targets were defined from the extracted HiChIP anchors via ChromHMM annotation. Next, the trained model and the functional genomics data of interest get put through anchor type predictor (ATP), which then identifies the putative topological interactions existing between active genomic regions. The anchor type output from ATP, the newly generated features, and targets of anchor pairs (enhancers and promoters) then get imported into our Confidence Predictor (CP) following gradient boosted regression trees (GBRT)-based training. Finally, CP assigns a confidence metric to each predicted chromatin loops, which can be utilized for the filtering of the final LoopPredictor output. (B) Distribution of general HiChIP loop scores after merging four HiChIP datasets (K562-H2K27ac, K562-YY1, HCT116-YY1, and GM12878-H3K27ac). (C) Diagram depicting the loop associated regions used to gather features in Confidence Predictor (CP).

The first component of the algorithm core is ATP, which is a minimal classifier based on Random Forest (27) and multi-task frameworks that uses a minimum number of features to get optimal prediction power, and then identifies the possible conformation between genomic regions. Four HiChIP datasets were processed by HiC-pro (28) and the loops were generated by hichipper (29) (FDR<0.05, and pair-end tag number ≥2). Next, we extracted the anchors from loops, and used ChromHMM (30) to annotate the chromatin state of each anchor. The annotations of anchors were regarded as targets. We collected a variety of multi-omics data for the feature generator, which used a standard scaler for normalization and batch effect removal. A hybrid Random Forest was applied to train the model utilizing the multi-task framework. After training, datasets of interest are used as input (e.g. H3K27ac ChIP-seq peaks) into the ATP model to generate the possible anchor pairs and predict the type of anchors for loop prediction.

The second component of the algorithm core is CP, which is a powerful regressor based on Gradient Boosted Regression Trees (GBRT). The possible conformation and the corresponding loop type generated by ATP were imported into CP to predict the confidence level as a loop score. We found that the scores of loops obey a gamma distribution (**Fig. 2B**), so we used the density function to identify high confidence loops and then normalized the scores for target generator. Next, we integrated more features for CP from different genomic scales, including the flanking regions of two anchors, the distance between two anchors (window), and the neighboring regions outside of each anchor (**Fig. 2C**). The anchor type from ATP, and the newly generated features and targets of anchor pairs were imported into GBRT for training. Finally, we used CP to predict chromatin loops with scores indicating the confidence of the topological interaction.

### The performance of Anchor Type Predictor

Before determining which classifier to be used in ATP, we tested the F1-score of four standard classifiers: LinearSVC (31), LogisticRegression (31,32), KNeighbors (31), and Random Forest (31,33). The F1-score is the harmonic mean of the precision and recall, which is often used to evaluate the performance of machine learning models. The classifiers were trained with four different HiChIP datasets (K562-YY1, K562-H3K27ac, HCT116-YY1, and GM12878-H3K27ac). The evaluation of the F1-score showed that Random Forest (RF) outperforms the other methods (**Fig. 3A**). Similarly, the Receiver Operating Characteristic (ROC) curves (34) of these four classifiers in K562-YY1 HiChIP data indicated that RF achieved the best in the prediction of different kinds loops (promoter-enhancer, promoter-promoter, enhancer-enhancer) (**Fig. 3B**). Therefore, we integrated RF into ATP to generate the possible anchor pairs and predict the type of anchors.

**Figure 3.**
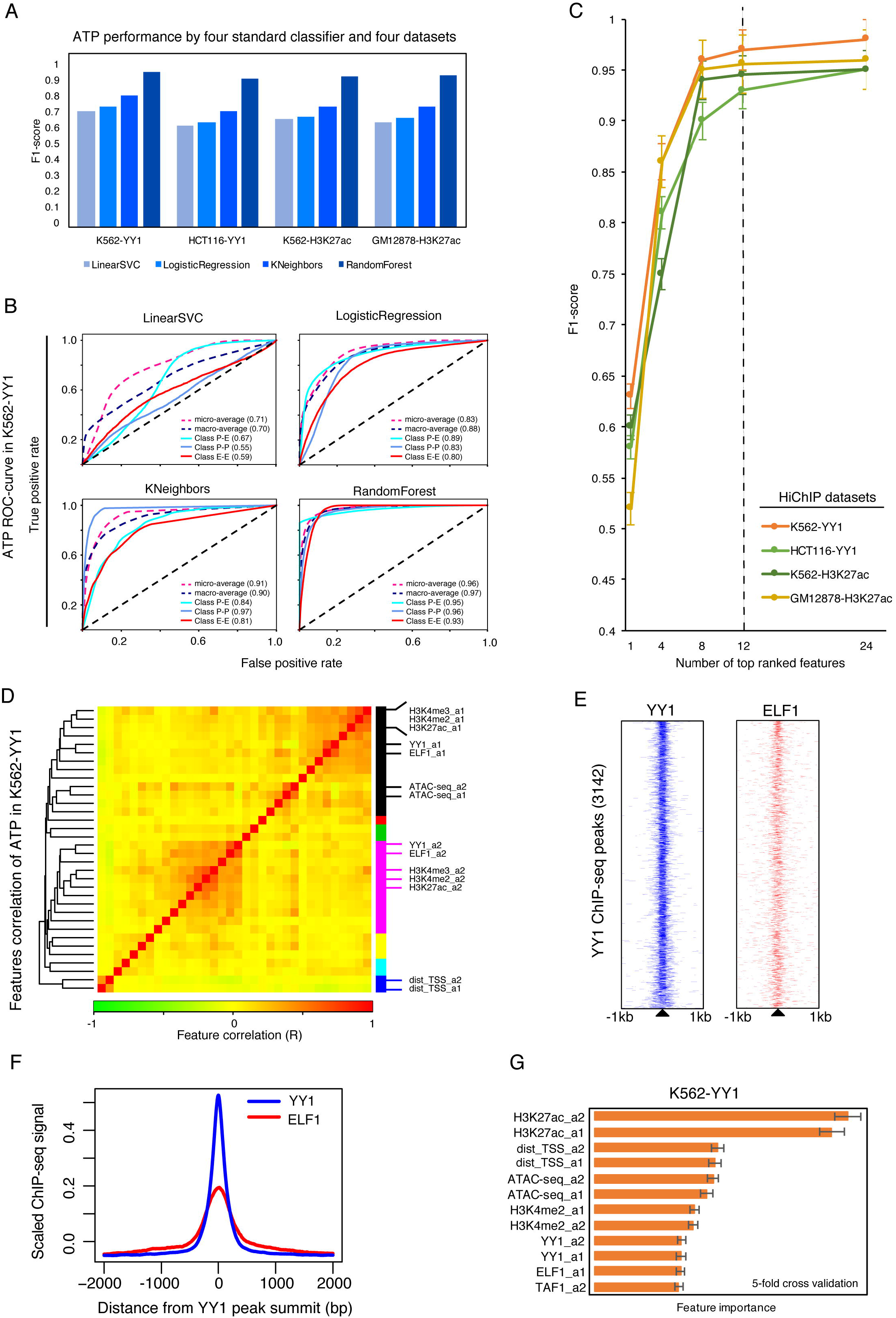
The performance of Anchor Type Predictor (ATP). (A) F1-score of ATP evaluated using four standard classifiers across four HiChIP datasets. (B) ROC curves of ATP in K562-YY1 HiChIP data. (C) The F1-score performance of ATP in four HiChIP datasets with increasing number of top ranked features by 5-fold cross validation. The vertical line indicates point at which ATP achieved close to optimal performance with a modest input of 12 features. (D) Pearson correlation combined with hierarchical clustering for K562 cell features. The colored bars on the right side indicate the identity of each hierarchical cluster. Colored bars at the bottom mark the feature correlation coefficient, R. (E) Heatmap displaying YY1 and ELF1 ChIP-seq signals across YY1 peaks (n = 3,142). (F) Comparison of YY1 and ELF1 ChIP-seq peaks signals by distance from YY1 peak summit. (G) Feature importance for the top 12 features in K562-YY1 HiChIP dataset with 5-fold cross validation. Error bars indicate standard deviation.

To obtain the minimum input for optimal performance of ATP, we tested the F1-score with an increasing number of features (total n=24). A feature is a multi-omics dataset (e.g. H3K4me1 ChIP-seq). The performance with 12 features was close to optimal, so we chose this number of features to use as input for RF (**Fig. 3C**). To determine the correlation between features, Pearson correlation and hierarchical clustering analysis were used for all 24 features (**Fig. 3D**). Importantly, active histone marks, ATAC-seq, and YY1 ChIP-seq signals were highly correlated. Moreover, we found that ELF1 ChIP-seq was also highly correlated with YY1. ELF1 is a lymphoid transcription factor known to regulate the expression of MEIS1 (in K562 cells), another transcriptional master-regulator associated with leukemic hematopoiesis whose motif was found to be enriched in H3K27ac anchors (35). Analysis of YY1 and ELF1 ChIP-seq data confirmed the co-localization of the two factors (**Fig. 3E**, and **3F**). Thus, the correlations uncovered here are likely to be functionally relevant. The feature correlation of K562-H3K27ac, HCT116-YY1, and GM12878-H3K27ac HiChIP datasets were also similarly clustered (SI Appendix, **Fig. S1**). We then investigated the importance of our top 12 ranking features from the K562-YY1 HiChIP dataset using a 5-fold cross validation procedure. The data showed that the H3K27ac ChIP-seq signal within two anchors was the most important feature followed by the distance from anchors to transcription start sites (TSS) and chromatin accessibility (**Fig. 3G**). The feature importance of other HiChIP datasets was similar to K562-YY1HiChIP (SI Appendix, **Fig. S1**).

### The performance of Confidence Predictor (CP)

To characterize the general profile of all the features in different loop-associated regions (two outer neighbors, inter-anchor window, and two anchors), we quantified each individual feature signal on a z-score normalized scale (36). The feature signal of the inter-anchor window region was highest and most variable, while the outer neighbor regions presented the lowest intensity (**Fig. 4A**). Here we trained CP with four individual HiChIP datasets (K562-YY1, HCT116-YY1, K562-H3K27ac and GM12878-H3K27ac) and four integrated datasets, including K562(YY1+H3K27ac)*, K562*+HCT116, K562*+GM12878, and K562*+GM12878+HCT116, the star symbol was used to present the combination of K562-YY1 and K562-H3K27ac, as these two datasets are from the same cell line, so we combined them for the integration. To interpret the contribution and correlation of features in the prediction, we calculated an importance score for each feature with 5-fold cross validation, and then filtered the features whose importance score was greater than 0.001 for Pearson correlation analysis and hierarchical clustering (**Fig. 4B**, and SI Appendix, **Fig. S2**). The data showed that features are correlated well by loop-associated regions, and the size of window was the most important factor for the prediction, which was consistent across the cell lines analyzed.

**Figure 4.**
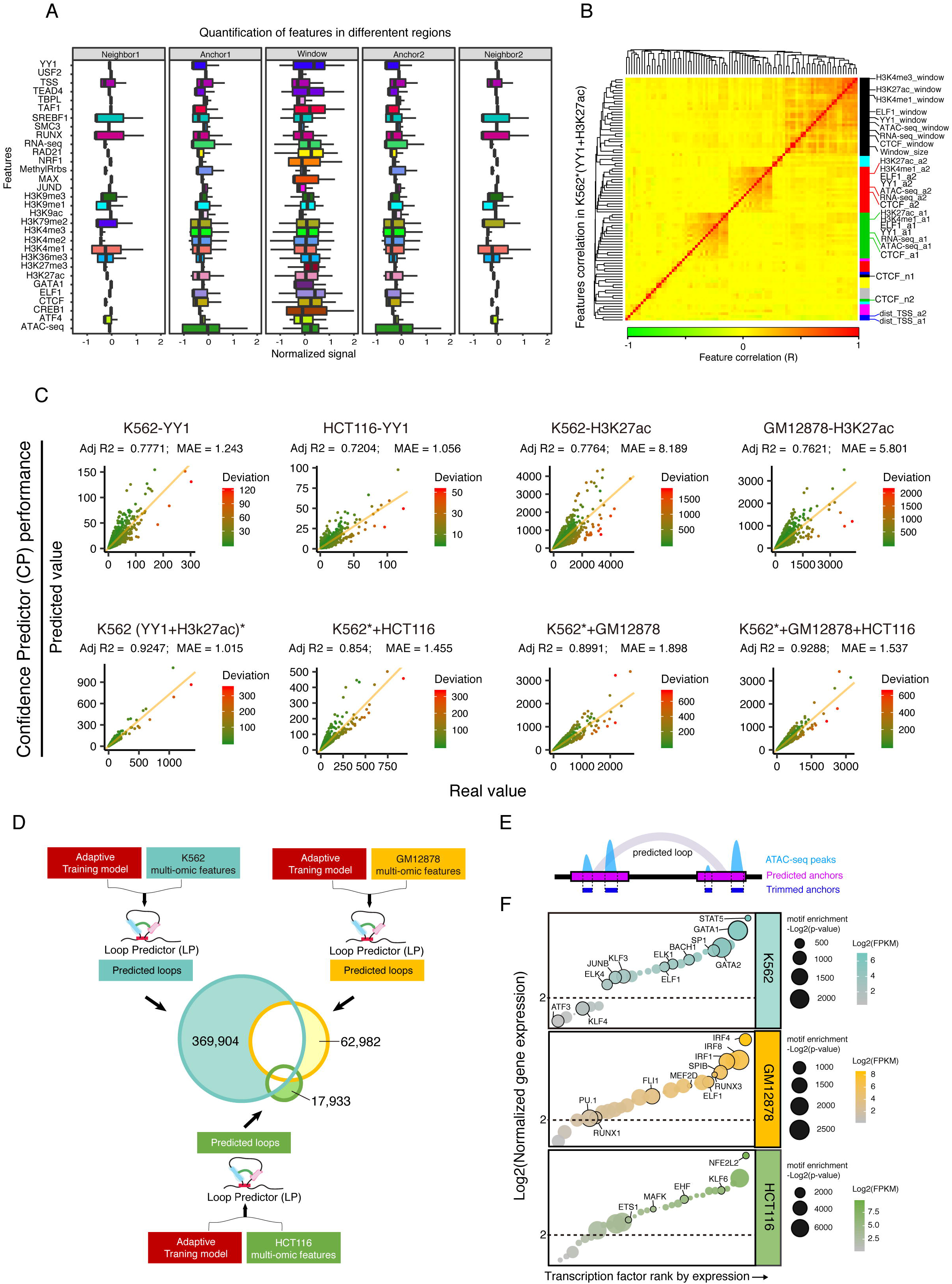
The performance of Confidence Predictor (CP). (A) Quantification of features across different loop associated regions. The signals of features were normalized by z-score. (B) Pearson correlation and hierarchical clustering for the features of K562*(YY1+H3K27ac). The colored bars on the right side indicate hierarchical clusters. Colored bars at the bottom demarcate the feature correlation coefficient, R. (C) Prediction performance evaluation for CP in four individual datasets (upper) and four integrated datasets (below). (D) Evaluation of LoopPredictor for identifying cell type-specific loops. Our adaptive model and the multi-omics features derived from three individual cell lines were fed into LoopPredictor to perform predictions. The results from each cell type were marked by different colors in the Venn diagram. The number of predicted cell type specific (K562, GM12878, HCT116) loops were 369904, 62982, and 17933, respectively. (E) Diagram for trimming anchors with ATAC-seq peaks to identify loop binding transcription factors. (F) Transcription factor enrichment analysis for cell type-specific predicted loops. Cell type-specific loops identified in Fig. 4D. Transcription factors were ranked by normalized gene expression. The size of each point indicates the motif enrichment *P* value. The color of each point codes for the normalized expression of the indicated transcription factor. The threshold of motif enrichment was −log2(p-value) > 500, and the threshold for gene expression was set to log2(FPKM) > 2.

To evaluate the performance of CP, we calculated the adjusted R-square value and Mean Absolute Error (MAE) for different prediction cases, and then assessed actual values versus predicted observations (**Fig. 4C**). For the prediction of four individual dataset (K562-YY1, HCT116-YY1, K562-H3K27ac and GM12878-H3K27ac), CP achieved an adjusted R-square from 0.72 to 0.77, while for the integrated datasets, the adjusted R-square values of CP were all larger than 0.85. Importantly, the integration of K562* (YY1+ H3K27ac), GM12878, and HCT116 outperformed the others with the highest adjusted R-square of 0.9288 with an MAE of 1.537 (**Fig. 4C**). Specifically, the distribution of actual loop scores and predicted loop scores are consistent (SI Appendix, **Fig. S3A-S3D**). These results suggest that the integrated datasets are more favorable for the training of CP.

We next assembled the ATP and CP modules together into an adaptable model, which were trained with the HiChIP integrated datasets. Next, the adaptable model and multi-omics features from different cell types were fed into LoopPredictor to predict enhancer-mediated interactions. One concern with utilizing our multicellular multi-omics trained adaptive model is a loss of cell-type specific loops, and a potential enrichment for common regulatory genomic interactions. To evaluate the performance of our adaptive training model for predicting cell type-specific observations, we fed multi-omics features from three different cell lines separately (**Fig. 4D**). Importantly, we identified thousands of unique loops for each input cell line. To determine the regulatory characteristics of these unique loops, we extracted the predicted anchors and overlapped them with cell line-specific accessible chromatin peaks (ATAC-seq and DNase-seq) (**Fig. 4E**). The highest enriched motifs from these three trimmed anchor sets were extracted and ranked by gene expression to produce a list of cell line-specific transcription factors (**Fig. 4F**). Indeed, we identified GATA1 activity in K562 cells, enrichment for IRF factors in GM1212878 loops, and NRF2 binding in HCT116 cells, consistent with the literature (37–39). Hence, our comprehensive adaptive training model is a powerful tool for predicting cell type-specific enhancer promoter loops.

### Functional validation of predicted enhancer-mediated interactions

To investigate the degree to which the predicted loops output from LoopPredictor matched with experimentally measured enhancer-promoter loops, we compared the predicted loops from K562 cells with H3K27ac HiChIP performed in K562 cells. The K562 predicted interactions generated by LoopPredictor contained 789,297 loops with a confidence score threshold of 7 covered 95.9% of the K562-H3K27ac HiChIP loops, while only 4.1% (12,922) loops were K562-H3K27ac HiChIP specific (**Fig. 5A**). To investigate the distribution of loops by distance, we binned the loops by 100 kb windows. The resulting loop proportions were consistent between the predicted universal interactions and the observed H3K27ac HiChIP interactions (**Fig. 5B**). Differential analysis of loop scores also indicated that most loops were not significantly different between K562 predicted and K562-H3K27ac HiChIP loops (**Fig. 5C**).

**Figure 5.**
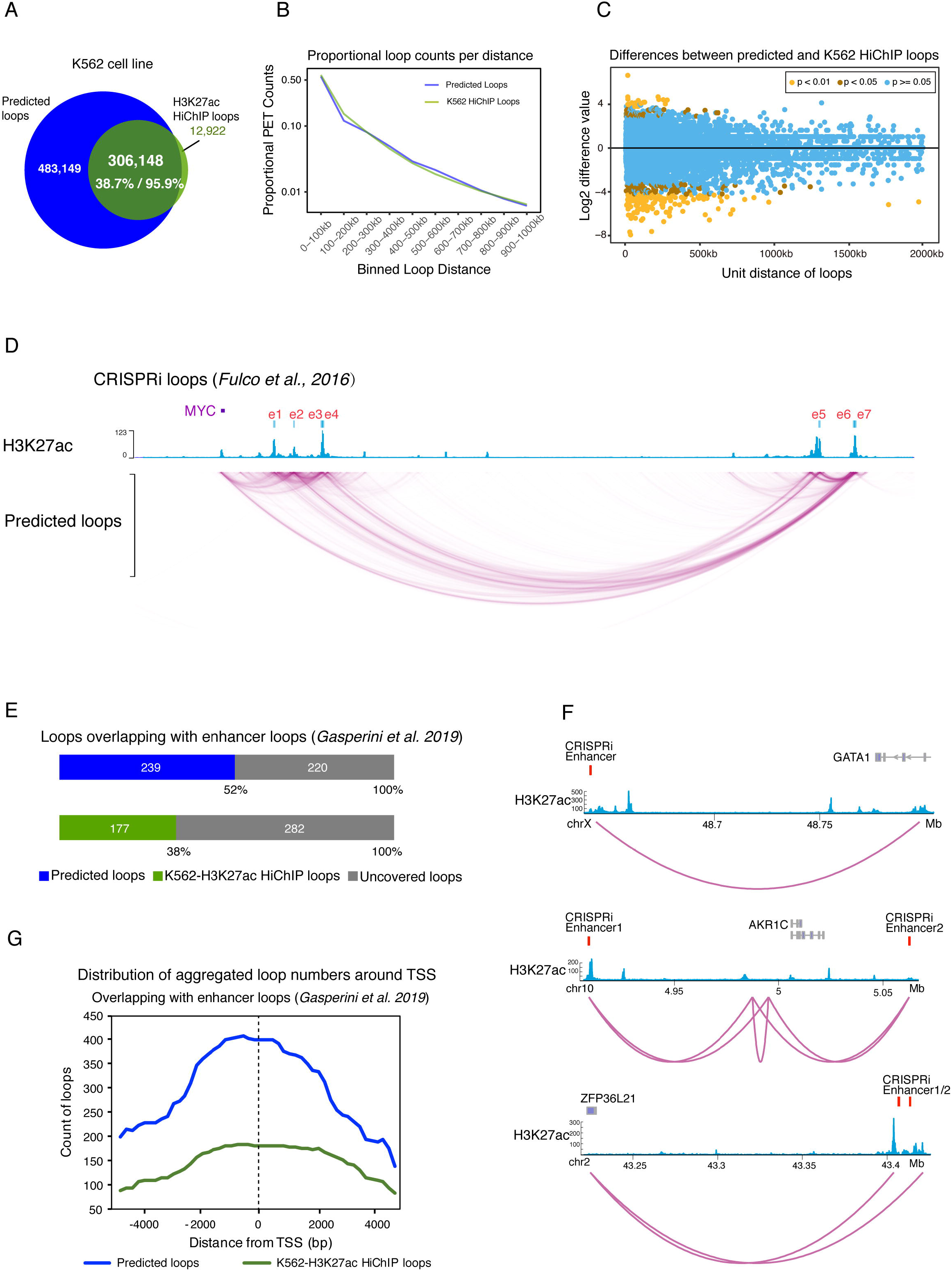
Functional validation of predicted enhancer-mediated interactions. (A) Venn diagram showing the overlap of K562 predicted loops and K562-H3K27ac HiChIP loops. Overall, 306,148 loops were detected by both LoopPredictor and HiChIP experiments, which accounted for 95.9% of the HiChIP loops. 4.1% (12922 loops) were HiChIP-specific loops. (B) Proportional loop counts per distance for predicted and K562-H3K27ac HiChIP loops. Loops were binned by 100 kb to calculate the proportion. (C) Differences between predicted and K562-H3K27ac HiChIP loops. Loops with a p-value >= 0.05 (blue dots) were classified as non-significant, loops with a p-value < 0.05 (brown dots) were labelled significant, and differences with a p-value < 0.01 (yellow dots) were marked highly significant. The vast majority of loops of loops showed no significant differences between the two sets of loops. (D) Validation of predicted loops by focused CRISPRi integration (*Fulco et al. 2016*). Seven previously validated *MYC* enhancers with strong H3K27ac ChIP-seq signals (blue track) were annotated as e1 through e7 (red track). The predicted loops contacted these published CRISPRi loops. (E) Validation of predicted loops by high-throughput CRISPRi screening integration (*Gasperini et al. 2019*). A total of 459 high-confidence gene-enhancer loop pairs were overlapped with predicted loops as well as H3K27ac loops. In total, 52% of the high-confidence loop pairs were identified by LoopPredictor. Only 38% of these high-confidence loop pairs were recovered from K562-H3K27ac HiChIP loops. (F) Genome browser tracks for validated enhancer loops identified in Fig. 5E. Validated CRISPRi high-confidence pairs are shown in red. H3K27ac ChIP-seq (blue). Predicted loops (purple). (G) Promoter contacts of CRISPRi loops for predicted loops, and H3K27ac HiChIP loops. Distribution of aggregated loop numbers by distance of loops from Fig. 5E. The distribution was calculated by ± 4 kb distance from TSS.

We next wanted to validate our predicted loops by comparing them to functionally validated enhancer-promoter pairs identified in K562 cells. Previously, 7 *MYC* enhancers were identified via a systematic CRISPR interference (CRISPRi) screen, which were annotated as e1 through e7 (40). Notably, the predicted loops output from LoopPredictor proximal to *MYC* were in accordance with these published loops: MYC-e1, MYC-e2, MYC-e3/e4, MYC-e5, MYC-e6/e7, e5-e6/e7 (**Fig. 5D**), indicating a high accuracy of predictive power. Recently, 664 enhancer-gene loops were identified from a large-scale multiplex enhancer-gene pair screening effort in K562 cells (41). From this study, we identified the high-confidence enhancer-gene loops (n=470 pairs) for comparison with our predicted K562 loops. Indeed, 52% of the functionally validated enhancer loops overlapped with our predicted loops, compared to just 38% overlap with K562-H3K27ac HiChIP loops (**Fig. 5E**, and **5F**) (10). Within these overlapping loops, the predicted observations had stronger enrichment of loop counts proximal to the TSS compared to H3K27ac HiChIP loops (**Fig. 5G**). Hence, LoopPredictor is capable of predicting functional enhancer-gene loops with high sensitivity.

### Predicting chromatin interactions in a model organism

LoopPredictor can predict the topological interactions for any cell types which lacks 3D genomic information, and because it’s trained on highly conserved mammalian gene regulatory features it should also be able to predict enhancer-promoter interactions for other mammalian species (e.g. mice) (42). We gathered multi-omics features from the murine NIH3T3 myofibroblasts cell line, and the aforementioned adaptable model trained on human cell lines to feed into LoopPredictor (**Fig. 6A**). After the prediction, we obtained 59,708 loops as output, the proportional loop counts by distance showed a high coincidence between the predicted loops and published NIH3T3 HiChIP loops (**Fig. 6B**) (18), and the differential analysis showed that most of the loops had no significant difference (SI Appendix, **Fig. S3E-S3G**). To interpret the component differences between predicted loops and NIH3T3 HiChIP loops, we annotated the anchor types by ChIP-Seq data and the distance from anchors to TSS (**Fig. 6C**). The anchors were classified into three types: Enhancers (E), Promoters (P), and Other (O). Annotation results showed that enhancer anchors accounted for the majority of anchors in all loop sets. Interestingly, the overlapping HiChIP and predicted loop anchor were comprised entirely of enhancer and promoters, while there were 41.7% O-type loops in the NIH3T3-specific HiChIP loops, which was higher than the LoopPredictor-specific loops (6.4%) (**Fig. 6C**). We hypothesized that NIH3T3-specific non-enhancer-promoter anchors may lie within inactive heterochromatic regions, so we analyzed NIH3T3 H3K27me3 CUT&RUN data over these O-type anchors (**Fig. 6D**). NIH3T3 HiChIP-specific and LoopPredictor-specific O-type loops both displayed high H3K27me3 signals suggesting that these anchors primarily lie in inactive genomic regions. Thus, the loops output from LoopPredictor are primarily active loops with decreased heterochromatin composition compared to H3K27ac HiChIP-specific loops.

**Figure 6.**
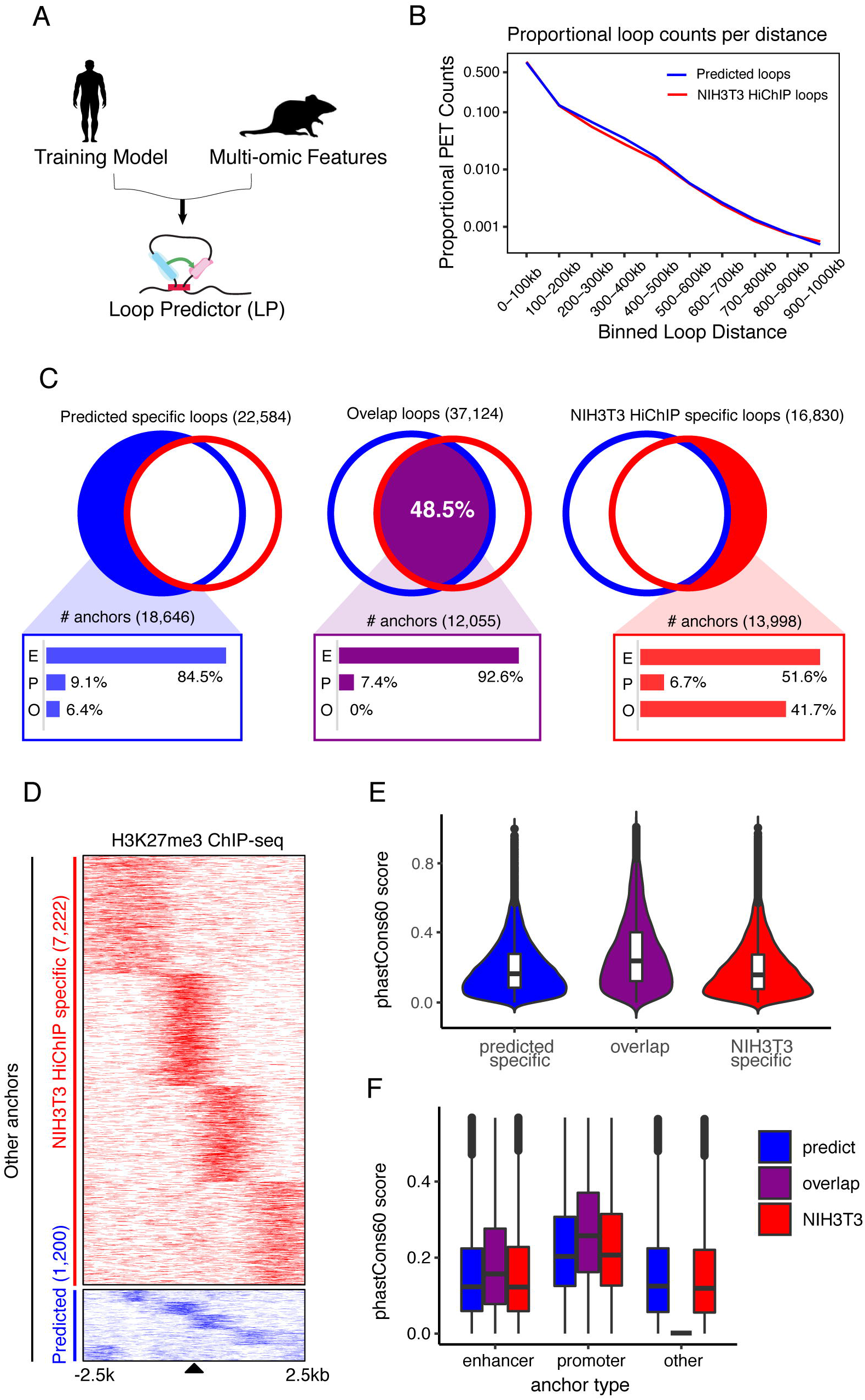
Cross-species long-range chromatin loop predictions. (A) Diagram for cross-species chromatin predictions. The adaptive training model implemented previously, and trained on human cell lines, was combined with multi-omics data from murine NIH3T3 myofibroblasts as input for LoopPredictor. (B) Proportional loop counts per distance for predicted and NIH3T3 HiChIP loops. Loops were binned by 100 kb to calculate the proportion. (C) Venn diagrams depicting the amounts of predicted-specific loops (22,584), overlapping loops (37,124) and NIH3T3-specific HiChIP loops (16,830). The common loops accounted for 48.5% of the total loops. The bar plots shown below indicate the composition of anchor types from each loop set. Enhancer type (E), Promoter type (P), and Other type (O). (D) Heatmap for H3K27me3 ChIP-seq signal over O-type anchors in NIH3T3 HiChIP-specific and predicted-specific loops found in Fig. 6C. The number of O-type anchors in NIH3T3 HiChIP-specific and predicted-specific sets are 7,222 and 1,200, respectively. ChIP-seq signals were visualized across a 5 kb window. (E) Violin plots of conservation scores across anchor types. Anchor regions derived from Fig. 6C. (F) Boxplot for conservation scores by anchor type derived from Fig. 6C.

As we conducted a cross-species comparison, we next investigated the degree of conservation between human and mouse over the LoopPredictor-specific, NIH3T3 HiChIP-specific, and overlapping loops (**Fig. 6E**). The mean conservation score of overlapping loop anchors was greatest among the 3 groups (**Fig. 6E**). Moreover, the promoter anchors were the most conserved anchor type (**Fig. 6F**). This is consistent with previous large-scale regulatory studies, which found that the binding of orthologous transcription factors to promoters is more highly conserved than binding to distal regulatory sequences (42). Thus, LoopPredictor is capable of performing cross-species loop predictions with improved sensitivity over running H3K27ac HiChIP alone.

## DISCUSSION

Enhancer-mediated interactions play important roles in gene expression, evolution, disease, and development. Currently, it is a significant challenge to investigate the genome topology for all cell types across species. Therefore, we generated LoopPredictor, an ensemble machine learning model to predict the genome topology for any cell type which lacks a 3D profile. LoopPredictor incorporates H3K27ac/YY1 HiChIP datasets and an assembly of multi-omics features to learn active long-range enhancer mediated looping characteristics. Users need only to provide multi-omics datasets as input, alongside our adaptive training model, into LoopPredictor to generate a list of predicted loops ranked by confidence, and comprehensively annotated.

The adaptive training model we incorporated for predicting loops was generated using many multi-omics datasets derived from several distinct cell lines. Despite the diverse cellular input underlying our model, we were able to identify thousands of cell type-specific enhancer loops with LoopPredictor when appropriate inputs were added. Indeed, we were able to isolate cell type-specific gene regulatory networks from among three different cancer cell lines. And in addition to the factors we identified that were consistent with what has previously been reported in the literature, we identified several more that likely have underappreciated roles in long-range enhancer mediated gene regulation. Moreover, the comprehensive list of cell type-specific transcription factors, along with their predicted binding sites, may be useful for the design of high throughput screens aimed at understanding processes, like oncogenesis, cell identity, metastasis, and basic mechanisms of gene regulation.

Existing computational methods to predict enhancer-promoter interactions often rely on a simple assumption that connects a single peak or region (e.g. ATAC-seq peak) to the nearest gene. More sophisticated computational methods, like TargetFinder, incorporate diverse multi-omics features to predict enhancer-promoter interactions with improved accuracy compared to using only the closest gene (43–45). Indeed, the predictive importance of genomics features output from TargetFinder were remarkably similar when directly compared with LoopPredictor. For example, H3K4me3, H3K4me1, H3K27ac, and DNase-seq signals were found to be high importance features within inter-anchor window regions for both methodologies. Interestingly, YY1 was identified by TargetFinder as having low to no importance within enhancers and no reported importance within promoters. However, within our model the feature importance of YY1 is most significant within the window region. Many possibilities can explain the differences in feature importance between the two pipelines, including differences in datasets incorporated, and overall algorithm design. One important distinction between TargetFinder and LoopPredictor is the use of HiC data as opposed to H3K27ac- and YY1-HiChIP data, respectively. The majority of loops identified from HiC data are thought to be large structural CTCF and cohesion anchored loops (9). However, the HiChIP loops incorporated into LoopPredictor are enriched for YY1 and H3K27ac bound regions and should thus have a higher proportion of active enhancer mediated loops, and subsequently a decreased amount of background CTCF-associated interactions (9). The differences in 3C-based technologies incorporated may explain why LoopPredictor’s near-optimal performance was achieved with 12 features, while TargetFinder required approximately 16. Further, our model offers several other advantages over existing predictive tools, including the ability to detect enhancer-enhancer, enhancer-promoter, and promoter-promoter interactions on a genome-wide scale. This utility may allow for the mapping of complex 3D enhancer cliques. And LoopPredictor can predict topological interactions for cell lines which lack any pre-existing 3D data, unlike TargetFinder which can only identify enhancer-promoter interactions for cell lines which have accompanying HiC data.

The ensemble model implemented by LoopPredictor consists of two core components, ATP, and CP. For the classification step performed by ATP, we tested performance with the F1-score for different cell lines and different classifiers, and the results indicated that Random Forest performed the best in all datasets. And the classification capability of ATP was optimal with an input of 24 features, while 12 features was recommended, which reduced the input by half. The ranking of feature importance indicated H3K27ac and YY1 were key regulators for enhancer-promoter loops, which contributed to the identification of possible conformation and the classification of loop types. The normalized quantification results of features at different regions indicated the inter-anchor window region was most informative, as mentioned above, while the signal derived from the outer anchor regions was lowest. Moreover, most of the features were correlated well by window or anchor regions, while the features of two neighbors were scattered with no obvious association. The enrichment for feature signal between anchors is somewhat to be expected since the genomic regions between an active regulatory loop may contain more active enhancers, bound transcription factors, and highly accessible gene promoters. In contrast, genomic regions which are outside of anchors will more likely contain heterochromatic regions and 3’ genic regions. Future studies aimed at dissecting the components of active enhancer-promoter loops could benefit from performing a similar analysis and assessing these regions individually.

LoopPredictor, like H3K27ac HiChIP, has the ability to identify functional 3D enhancer loops. Here we found that after training our adaptive model and inputting several multi-omics features derived from K562 cells LoopPredictor predicted loops that were highly consistent with a published set of H3K27ac HiChIP loops derived from K562 cells. From this predicted loop set, we found an overlap for distal regulatory interactions between the *MYC* locus and seven enhancers which have been previously validated via CRISPRi screening (40). Moreover, a high throughput gene-enhancer pair screen performed in K562 cells identified several hundred high-confidence enhancer pairs which overlapped more favorably with the predicted loops compared to the experimental H3K27ac HiChIP loops (10). An explanation for these differences may be that the predicted loops, which are the result of a set of several combined functional genomic features and HiChIP loops, may be more sensitive than HiChIP alone. Our findings suggest that the predictive power achieved through incorporating multi-omics data with HiChIP loops is able to overcome dropouts of enhancer interactions from HiChIP datasets which may due to technical shortcomings of the assay, like GC content, length of interaction, sequencing depth, or chromatin composition. Alternatively, the result may be attributable to basic probability given that there were a greater number of observed predicted loops than experimental HiChIP loops.

The multi-omics features comprising the adaptive training model are overall very highly conserved, thus their predictive importance and the performance of LoopPredictor should be similar between humans and other mammalian species. To test the performance of LoopPredictor in other species, we fed multi-omics features derived from mouse NIH3T3 cells into the human trained adaptive model to predict murine loops. The predicted loops were nearly 50% identical with experimental NIH3T3 HiChIP loops. Further, the conservation of these identical overlapping sets of loops was greater than all non-overlapping loops. Thus, cross-species analysis with LoopPredictor can effectively predict conserved enhancer connectomes. Interestingly, the non-overlapping HiChIP-specific and predicted loops-specific signals contained detectable percentages of regulatory elements that couldn’t be annotated as either enhancers or promoters, the O-type elements. We found that these less conserved O-type elements, contained large amounts of the heterochromatin mark H3K27me3. These regions may represent background noise, or they may be comprised of boundary elements specific to the unique 3D nucleome of each species. Further detailed analysis of the composition and conservation of these elements is certainly warranted to evaluate the utility of LoopPredictor for performing comparative genomic analysis.

The wide-spread availability of high-throughput, low-input, and low cost multi-omics profiling technologies (e.g. CUT&RUN, and ATAC-seq) has dramatically increased the number of cell type-specific functional genomics datasets. Hence, there is a burgeoning need for tools to predict meaningful distal regulatory features with cell type-specific accuracy. LoopPredictor makes it theoretically possible to predict active regulatory topologies with high accuracy and sensitivity in all cell types which lack topological data.

## METHODS

### Identification of loops from HiChIP data

We collected four HiChIP datasets: K562-YY1, HCT116-YY1, K562-H3K27ac, and GM12878-H3K27ac for prediction, the raw reads in fastq format were downloaded from GEO. We employed HiC-Pro (28) to align the paired-end reads and used hichipper (29) to produce loops. Briefly, in the pipeline of HiC-Pro, the raw fastq HiChIP data were aligned to hg19 reference with “--very-sensitive” and “end-to-end” option. Then the alignment results as well as the restriction enzyme cut sites file (hg19 Mbol digest) were imported into hichipper, the chromatin loops were called by combining all the replicates and using all read density. We only considered the uniquely mapped reads with false discovery rate (FDR) < 0.05, the loops with at least 2 PETs supported and length larger than 5kb were retained. The vast majority of the loops (91.7% for K562-YY1,90.3% for HCT116-YY1, 96.3% for K562-H3K27ac, 95.9% for GM12878-H3K27ac) met above conditions and used for subsequent analysis.

### Identification of regulatory elements for all the anchors

Firstly, we extracted anchors from loops, then we utilized ChromHMM (30) annotations to identify the promoters and enhancers for the anchors from K562 and GM12878, and ENCODE Segway (46) for the anchors from HCT116 cell type. We also checked the transcribed activity of promoters by using the RNA-seq expression data from ENCODE (3), if the FPKM> 0.5, then retained the promoter.

### Super enhancer analysis

We employed the “-style super” option of findPeaks function in HOMER package (47) to identify the super enhancer regions, the peaks found within 12.5 kb were merged together into large regions, the slope threshold was set to 1000.

### GO analysis for enhancer anchors

GO analysis for enhancer anchors was taken by Metascape (48) with the minimum overlap of 3, and minimum enrichment of 1.5.

### Motif analysis for enhancer anchors

Firstly, the anchor regions from loops were overlapped and trimmed with the corresponding ATAC-seq peaks, then the trimmed anchors were used to detect motifs by HOMER package (47) with size of 200, the transcription factors with −log2(p-value)> 100 were chosen. Then the RNA-seq expression data from ENCODE was scaled by FPKM, the chosen transcription factors were ranked by the gene expression.

### Coincidence between different loop sets

We first used 500bp as a threshold to merge the nearby anchors into a valid anchor, if both of the valid anchors of one loop appear in another loop set, and there was an interaction between them, the loop is regarded as coincidence.

### Multi-omics datasets processing

Multi-omics datasets were prepared for the prediction, including ChIP-seq/CUT&RUN, ATAC-seq, in functional genomics, RRBS in epigenomics, and RNA-seq in transcriptomics. For ChIP-seq/CUT&RUN data, the peak files were downloaded from ENCODE if accessible, or used Bowtie2 to align the raw reads to reference with default settings, and uniquely mapping reads were used to identify enrichment regions. The sequence alignment was then transformed into platform independent data structure by makeTagdirectory package of HOMER, and findPeaks package was used to detect peaks with False discovery rate (FDR) <0.001, for transcription factor datasets, the peak size was set to 200bp, for histone marker datasets, the peak size was set to 500bp. The chromatin accessibility was profiled by ATAC-seq data, we filtered out some uninformative reads after the alignment with mapQuality<30, and “isProperPair” was set to only retain the proper paired reads, then the reads aligned to chrM were removed, MACS2 (49) was used to call peaks. The methylation profile of RRBS data were downloaded from ENCODE. The gene expression data were downloaded from ENCODE, or aligned the raw reads to reference by using STAR (50) with 2 mismatches at most, the raw read counts were normalized by FPKM.

### Features generation

For anchor type predictor (ATP), which is a minimal classifier, we trained the model with as few features as possible while ensuring the accuracy near optimal. Here, we only generated the features for the anchor regions. The R package GenomicRanges (51) was used to extract the profile values of specific regions. For ChIP-seq/CUT&RUN and ATAC-seq datasets, the peaks within anchor regions were extracted, and the mean value of peak signals were calculated as functional genomics features. For RRBS data, we calculated the methylation signals by multiplying methylation percentage by read counts for each profiled position within the anchor regions. The weighted mean values of methylation signals were used as epigenomics features. For RNA-seq data, we extracted the FPKM value of genes whose transcription start site locate within the anchors, then the mean FPKM value were used as transcriptomics features.

For Confidence predictor (CP), we constructed a powerful regressor to predict the score of loops accurately which required more features as input. We added left-flanking, in-between, right-flanking regions for each anchor pair to generate features, so there are five regions waiting for the feature generation including two anchor regions. The left-flanking regions were the 2kb extension from the start site of left anchor, the in-between regions were the intermediate parts of anchor pairs, the right-flanking regions were the 5kb extension from the end site of right anchor. The features generation method for anchor regions is consistent with ATP. For left-flanking, in-between, and right-flanking regions, we calculated mean values as well as standard deviations for every region following the same method for ATP.

### Training sample preparation

Four loop sets identified from HiChIP data were used as positive samples, the feature of each sample was generated as described above, and the annotations of regulatory elements for anchors were used as the target of samples, we only retained four types of targets for the prediction: promoter-enhancer, promoter-promoter, enhancer-enhancer, and none. The type of promoter-enhancer indicated one of the two anchors is promoter, and the other is enhancer, promoter-promoter and enhancer-enhancer indicated both of the anchors are promoters or enhancers. The type of none-none represented the loops are informative, including either of two anchors or both anchors are non-regulatory elements. Negative samples were produced by randomly selecting chromatin regions, avoiding ±2kb regions around TSS of any gene. The targets of negative samples were none-none, and the amount of negative sample was consistent with positive sample.

We combined positive and negative samples, and split the samples into 7:3 for training and testing, 5-fold cross validation was used in every training process.

### Classifier selection for anchor type predictor (ATP)

We tested the F1 scores of four standard classifiers: LinearSVC, LogisticRegression, KNeighbors, and RandomForest in four HiChIP datasets, four classifiers were constructed by using scikit-learn (31) with default parameters. The testing results is shown in Fig. **3A**, RandomForest outperformed the other classifiers, which was selected for the construction of ATP.

### A hybrid Random Forest classifier based on multi-task framework

Random Forest is a powerful classifier which uses all the given features to perform the prediction. As we need to train different classifiers for different HiChIP datasets, which is time consuming and tedious to select the important features to feed into model. Therefore, what we faced is how to select the most important features to minimize the input set while ensure the accuracy of prediction.

Multi-task learning is an approach which allows tasks training in parallel, and transforms information between related tasks, the inductive transformation would help each task learning better (52). Group Lasso is one of the sparse learning approaches, which utilizes the coefficients of features to construct the prediction model (53). In this study, we combined the feature selection ability of Group Lasso and the prediction power of Random Forest to construct a hybrid classifier. Then we built the hybrid Random Forest classifier on the framework of multi-task. Firstly, Group Lasso was used to explore the sparsity constraints of prediction, we defined the general classification task as *V_i_* = *α*_*i*_*m*_*i*_, *i* represents the number of sub-tasks, for the *i*-th task, *V_i_* is the labels vector for the task, and *α_i_* is the regression weights for *i*-th task, *m_i_* is the feature vector of *i*-th task, so we could use *M* to represent the feature matrix, in which *m_i_* is the *i*-th column, we assume there are N sub-tasks in total, the objective function is defined as

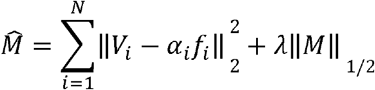

We applied the feature selection module before fitting Random Forest, the multi-task framework was implemented by scikit-learn. Then the hybrid classifier was integrated in ATP, we tested the performance of ATP in four HiChIP datasets by 5-fold cross validation, the importance of features was ranked for the prediction, which shows in Fig. 3E, and the F1 score performance of increasing feature numbers in Fig. 3C shows that ATP only needs 12 features as input to obtain near optimal.

### Features correlation evaluation

Firstly, we used “Pearson” method to calculate the correlation-based distance matrix for all the features, then applied hierarchical clustering on the matrix, which used the method of “average”. The correlation heatmap was implemented by using R packages “stats” and “gplots”.

### ChIP-Seq peaks density for YY1 and ELF1

The bed file of YY1 ChIP-seq peaks were used as the target regions, and the alignment file of YY1 and ELF1 in bigwig/bam format were used to plot the density heatmap, which was implemented by R package Genomation (54).

### ROC curve evaluation for ATP

Except for F1-score, we also utilized Receiver Operating Characteristic (ROC) metric to evaluate the classification quality of ATP. Regular ROC curves are used to evaluate the binary classification output, while our problem has four class labels. We therefore used the extension setting “Multiclass” in scikit-learn (31) to draw the ROC curve for each label. In addition, two kinds of measures were used to evaluate the general classification: “micro-averaging” considers each kind of label in the indicator matrix as a binary classification problem; “macro-averaging” allocates equal weight to each label in the classification task.

### Quantification of features in different regions

We gathered features for CP according to different loop-associated regions: two neighbors, two anchors and inter-anchor window. Then binned the features by the distance of 100bp, and the mean value and standard deviation of each region were calculated. We utilized z-score normalization to scale the signal of features, and then plotted the distribution of each region using boxplot from ggplot2 package (55).

### An adaptable Gradient Boosted Regression Trees (GBRT) regressor

Gradient Boosted Regression Trees (GBRT) is a kind of inductively generated tree ensemble model, which trains a new tree against the negative gradient of loss function for each step. The motivation of GBRT is to combine multiple weak learners to generate a powerful regressor. The additive model of GBRT was built in greedy function (56).

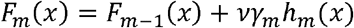

The tree newly added in each step was represented by *h_m_*, which tried to minimize the loss *L*, and GBRT used a type of negative gradient loos function for current model *F*_*m*−1_, *γ_m_* was step length, which was calculated by line search

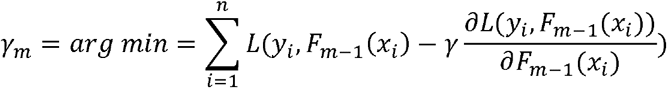

and *v* was used to scale step length, which called learning rate, learning rate impacted the training error cooperating with the number of weak learners. In addition, GBRT considered the strategy of stochastic gradient boosting (57), which combined gradient boosting with bagging, for each iteration, GBRT trained the base model on a fraction of training sample, and the value of fraction also impacted the performance of regression. Therefore, it’s crucial to determine the combination of learning rate, weak learner number and subsample fraction. Our problem is how to tune the model parameters for four different HiChIP datasets, meanwhile automatically adapt to the unknown datasets input by users. To solve the problem, we developed an adaptable module for GBRT to generate different combination of parameters to fit the model iteratively, then selected the optimal one to train the dataset and performed prediction.

### Evaluation of regression

We evaluated the performance of Confidence Predictor (CP) by calculating adjusted R-square value, Mean Absolute Error (MAE) and Root-Mean Squared Error (RMES). Adjusted R-square compares the explanatory power of regression models that contain different numbers of predictors, which is more objective than R-square to measure the multi-variable regression model. For R-square, the Sum of Squared Regression Error (RSS) and Sum of Squared Total Error were calculated, the calculation of adjusted R-square was based on R-square, which has been adjusted for the number of predictors in the model, and it is always lower than the R-square (58).

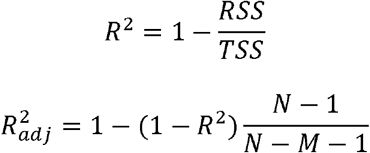

MAE measures the average magnitude of errors in a set of predictions without considering the direction. RMSE is a quadratic scoring function which measures the average magnitude of error by calculating the square root of the average between prediction and actual observation. The scatter plot of the predicted values versus the real values indicate the accuracy of prediction directly, the closer the slope of reference line to 1, the better the prediction.

### Validation of predicted loops

We used the same method with “Coincidence between different loop sets” to identify the overlapping loops. The proportion of loop counts by distance was calculated by R package diffloop (59), and the differential analysis between loop sets was implemented by R package HiCcompare (60). The visualization of H3K27ac ChIP-seq track and interactions was implemented by WashU Epigenome Browser (61). For calculating the distribution of aggregated loop numbers around TSS, we first annotated the anchor regions by ChIPpeakAnno (62), the distances between anchors and TSS were retrieved, and binned the loops by distances, then the number of loops for each bin was counted. The conformation plots of predicted loops were generated by Sushi (63).

### Conservation level evaluation for loops

The anchor regions of loops were overlapped and trimmed by the corresponding ATA-seq peaks, then the phastCons60 scores for trimmed anchors were extracted by GenomicScores (64).

## Published Datasets Used in This Study

ChIP-seq/CUT&RUN datasets: GSE29611 (3), GSE35583 (19), GSE31755 (3), GSE127432 (3), GSE32465 (20), GSE30263 (21), GSE31477 (3), GSE96253 (3), GSE92075 (3), GSE135286 (18), GSE63255 (22).

ATAC-seq datasets: GSE108513 (23), GSE47753 (24), GSE101975 (25), GSE135286 (18).

RNA-seq datasets: GSE88473 (3), GSE90276 (3), GSE33480 (3), GSE72860 (26).

RRBS datasets: GSE27584 (3), GSE27584 (3), GSE27584 (3).

## Acknowledgements

This work was supported by grants from the National Natural Science Foundation of China under Grants (No. 61732009) [M.L.], the 111 Project (No.B18059) [M.L.], Hunan Provincial Science and Technology Program (2018WK4001) [M.L.], the National Institutes of Health (DE023177, HL127717, HL130804, HL118761 [J.F.M.]; F31HL136065 [M.C.H.]; Vivian L. Smith Foundation (J.F.M.), State of Texas funding (J.F.M.), and Foundation LeDucq Transatlantic Networks of Excellence in Cardiovascular Research (14CVD01) “Defining the genomic topology of atrial fibrillation” (J.F.M.).

## Author Contributions

Conceptualization, L.T., and M.C.H..; Methodology, L.T., and M.C.H..; Investigation L.T., M.C.H..; Writing – Original Draft, M.C.H., and L.T.; Writing – Review & Editing, L.T., M.C.H., M.L., Jun W., and J.F.M.; Funding Acquisition, M.L, J.F.M., and M.C.H.; Resources, J.F.M. and M.L; Supervision, M.L, JX W., and J.F.M.; Visualization, M.C.H., and L.T.; Data Curation, L.T., and M.C.H.

---

**Supplementary Figure S1. Feature correlation and feature importance for ATP.** The colored bars on the right side indicate hierarchical clusters. Colored bars at the bottom demarcate the feature correlation coefficient, R. Error bars indicate standard deviation. (A) K562-H3K27ac HiChIP dataset. (B) GM12878-H3K27ac HiChIP dataset. (C) HCT116-YY1 HiChIP dataset.

**Supplementary Figure S2. Feature correlation and feature importance for CP in four individual datasets and four integrated datasets.** he colored bars on the right side indicate hierarchical clusters. The red-green colored bars at the bottom demarcate the feature correlation coefficient, R. Error bars indicate standard deviation. (A) K562-YY1 HiChIP dataset. (B) HCT116-YY1 HiChIP dataset. (C) K562-H3K27ac HiChIP dataset. (D) GM12878-H3K27ac HiChIP dataset. (E) K562-(H3K27ac+YY1)* HiChIP dataset. (F) K562*+HCT116 integrated HiChIP dataset. (G) K562*+GM12878 integrated HiChIP dataset. (H) K562*+GM12878+HCT116 integrated HiChIP dataset.

**Supplementary Figure S3. Distributions of actual loop scores and predicted loop scores.** (A) K562-YY1 HiChIP dataset. (B) K562-H3K27ac HiChIP dataset. (C) GM12878-H3K27ac HiChIP dataset. (D) HCT116-YY1 HiChIP dataset.

**Supplementary Figure S4. Predicting chromatin interactions in a model organism.** (A) Differences between predicted and NIH3T3-H3K27ac HiChIP loops. Loops with a p-value >= 0.05 (blue dots) were classified as non-significant, loops with a p-value < 0.05 (brown dots) were labelled significant, and differences with a p-value < 0.01 (yellow dots) were marked highly significant. The vast majority of loops of loops showed no significant differences between the two sets of loops.

(B, C) Some examples for comparing predicted and NIH3T3 HiChIP loops.

