## Supplementary figures and images for "Predicting unrecognized enhancer-mediated genome topology by an ensemble machine learning model"

### Supplemental figure1

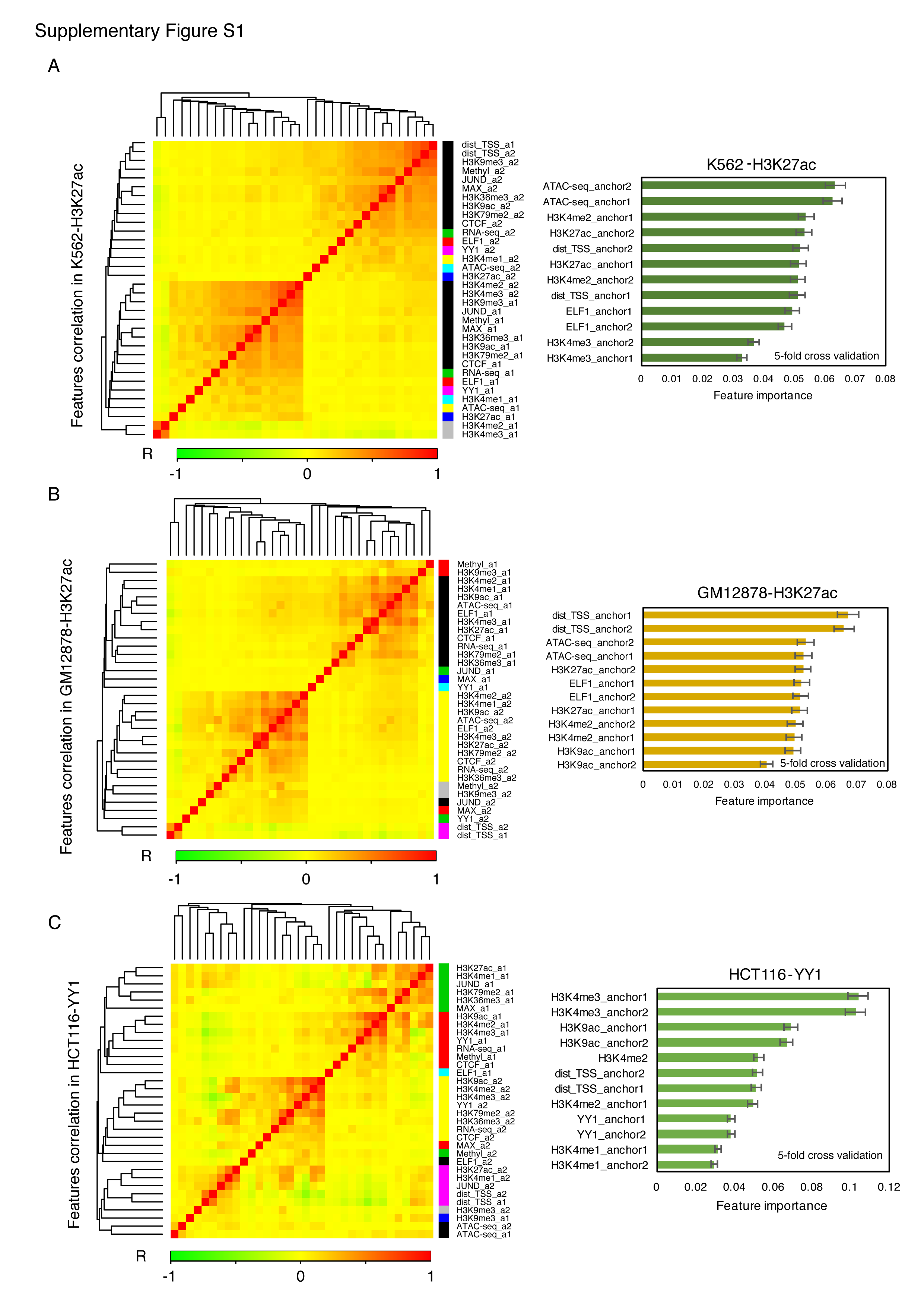

### Supplemental figure2

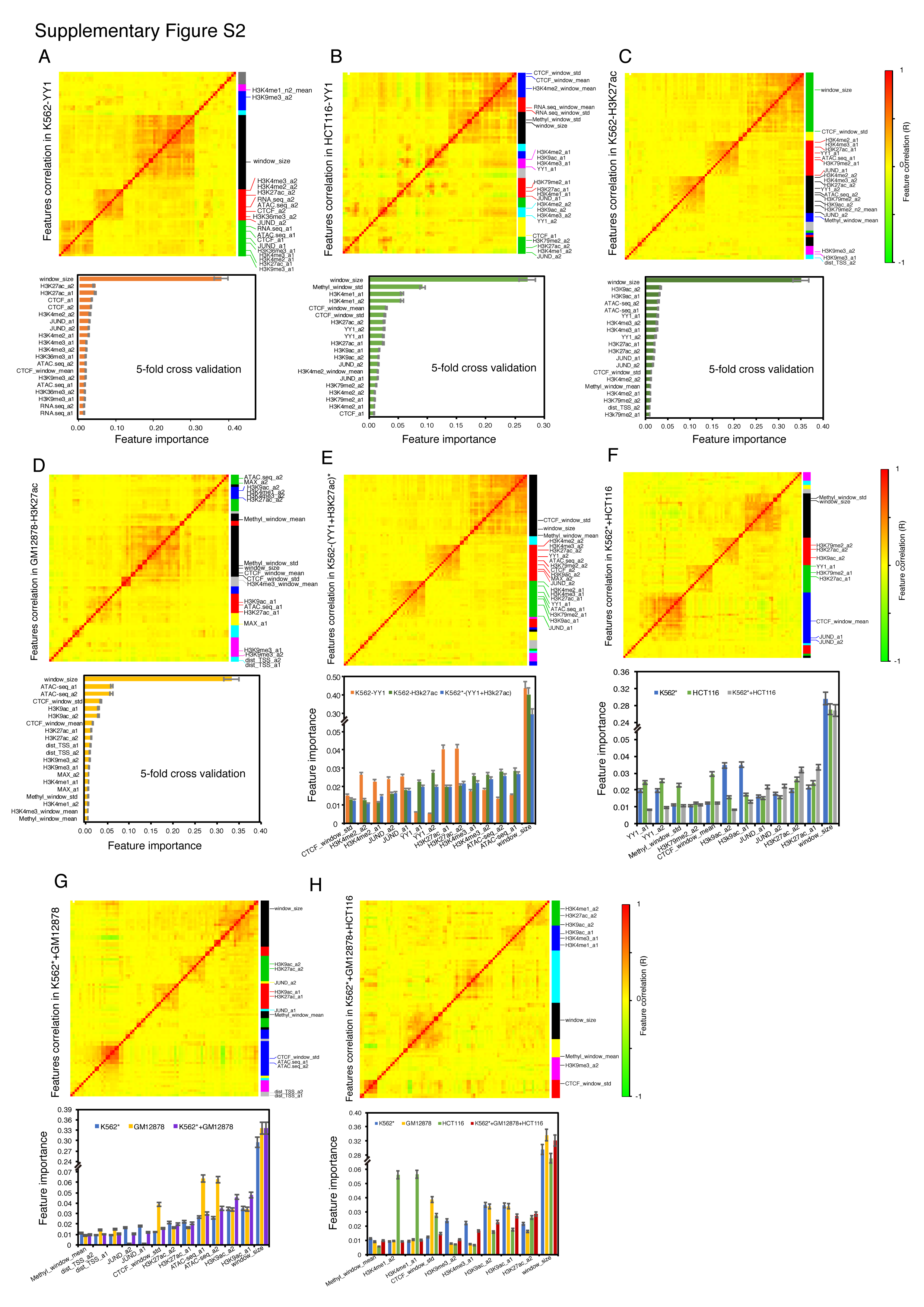

### Supplemental figure3

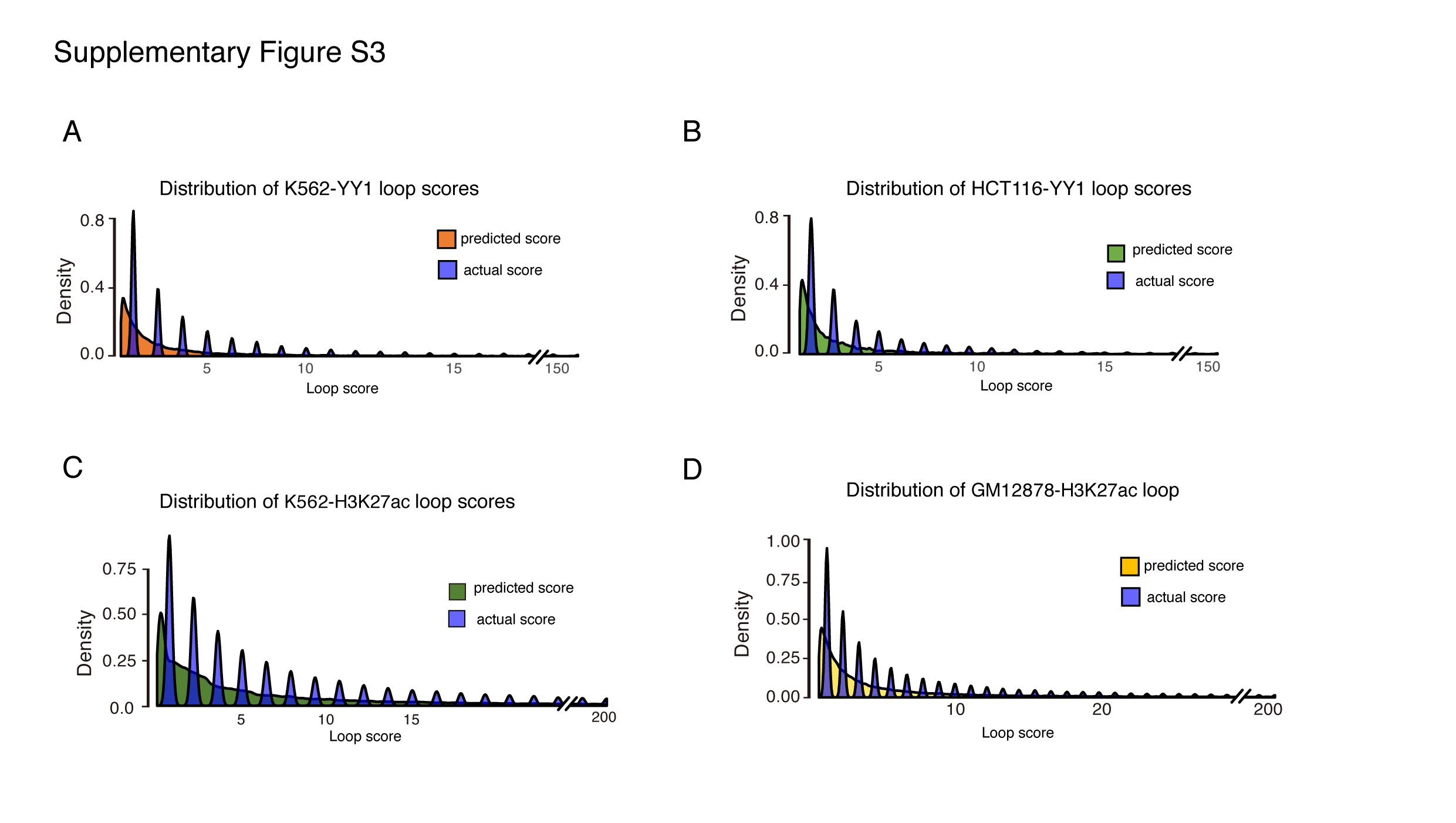

### Supplemental figure4

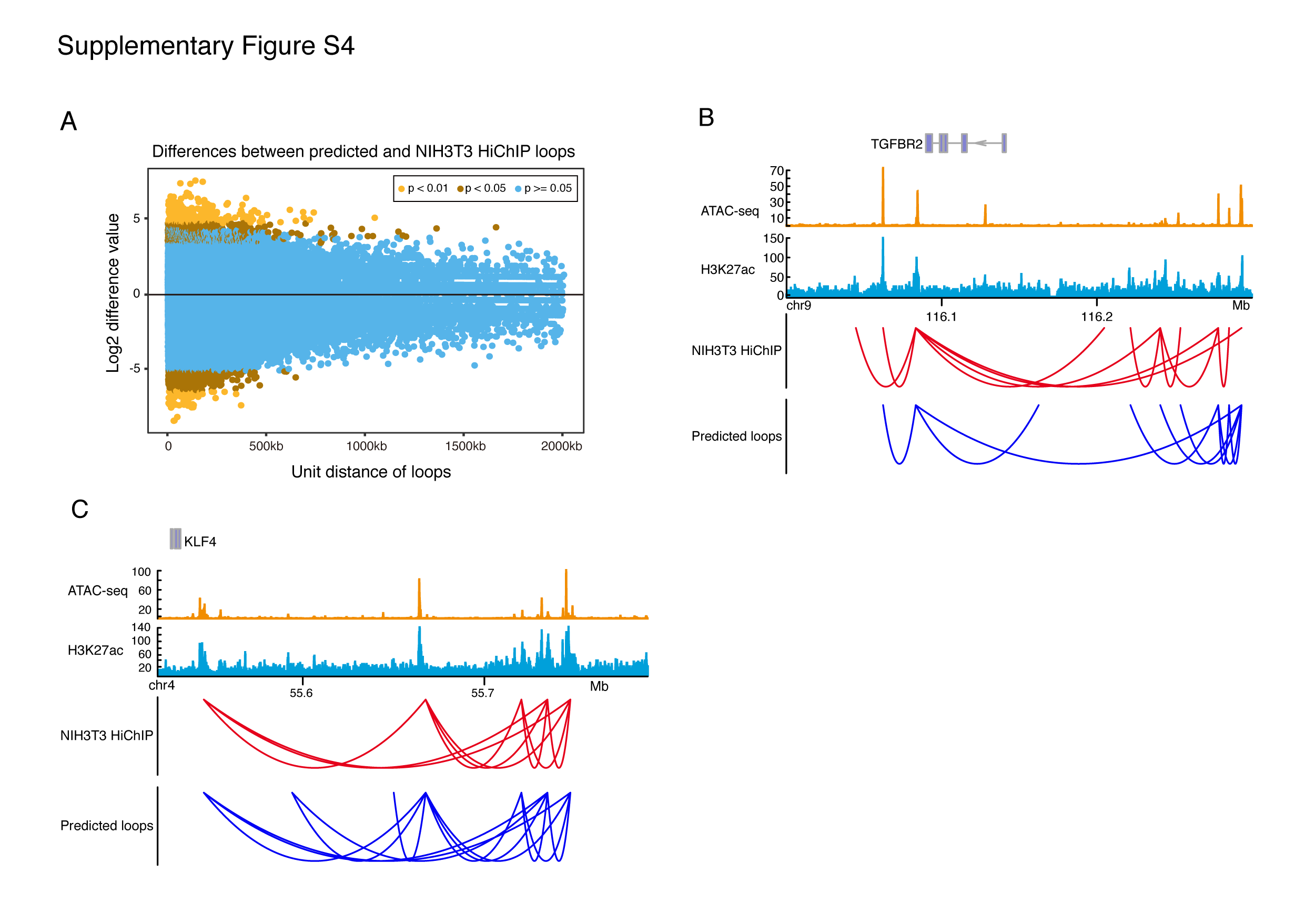
